# Electromechanics and Volume Dynamics in Non-excitable Tissue Cells

**DOI:** 10.1101/275339

**Authors:** F. Yellin, Y. Li, V. K. A. Sreenivasan, B. Farrell, M. B. Johny, D. Yue, S. X. Sun

## Abstract

Cell volume regulation is fundamentally important in phenomena such as cell growth, proliferation, tissue homeostasis and embryogenesis. How the cell size is set, maintained, and changed over a cell’s lifetime is not well understood. In this work we focus on how the volume of non-excitable tissue cells is coupled to the cell membrane electrical potential and the concentration of membrane-permeable ions in the cell environment. Specifically, we demonstrate that a sudden cell depolarization using the whole cell patch clamp results in a 30 percent increase in cell volume, while hyperpolarization results in a slight volume decrease. We find that cell volume can be partially controlled by changing the chloride or the sodium/potassium concentrations in the extracellular environment while maintaining a constant external osmotic pressure. Depletion of external chloride leads to a volume decrease in suspended HN31 cells. Introducing cells to a high potassium solution causes volume increase by up to 50%. Cell volume is also influenced by cortical tension: actin depolymerization leads to cell volume increase. We present an electrophysiology model of water dynamics driven by changes in membrane potential and in the concentration of permeable ions in the cell surrounding. The model quantitatively predicts that the cell volume is determined by the total amount of intracellular ion and protein content.

## 1 Introduction

Cells live in dynamic environments to which they must adapt [1, 2, 3]. In both physiological and pathological conditions, cells can respond to cytokines and other types of signals by changing their sizes [4, 5, 6, 7]. Cell volume changes can also trigger apoptosis, regulatory volume decrease, cell migration, and cell proliferation [8, 9, 10]. While it is well known that entropic forces from osmotic pressure differences can cause cell swelling or shrinking, changes in mechanical forces experienced by the cell can also influence cell volume [11]. For instance, active mechanical processes in the cell cytoskeleton, such as myosin contraction, generate contractile forces that impact cell volume regulation [12, 13]. Sudden changes in external hydrostatic pressure can change cell volume on the time scale of minutes [14]. Mathematical models of cell volume regulation have shown that there is a dynamic interplay between water flow, ionic fluxes, and active cytoskeleton contraction; all of these processes combine to influence cell mechanical behavior [15]. But many questions still remain: What are the factors determining homeostatic cell volume? How are cells able to sense volume changes? Moreover, cells live in saline environments where there are high concentrations of charged ions that are able to flow under electrical potential gradients. It has been shown that changing the transmembrane potential of non-excitable cells can affect cell shape, migration, proliferation, differentiation, and intercellular signaling [16, 17]. Since many of the same processes control both the cell osmotic pressure and membrane potential, we ask whether cell volume is closely coupled to membrane potential or the ionic environment. Indeed, cell volume changes have been observed when the ionic environment of the medium is modulated by applied electrical fields [18]. Previous experiments have explored shape changes in cells due to specific ionic currents or ion channels/pumps, e.g. the effects of Ca^2+^ on shape oscillations [19, 20] and regulatory volume decrease due to SWELL channels [21, 22, 23]. These studies do not treat the cell as an electro-chemo-mechanical system, but instead focus on specific signaling networks or ionic currents. In this paper, we aim to understand how mechanical, electrical, and chemical systems work together, with primary focus on the principal ionic components sodium (Na^+^), potassium (K^+^), and chloride (Cl^−^).

We first address whether the cell volume is related to the transmembrane electrical potential (Fig. 1). We perform whole cell patch clamp experiments [24] on suspended head-neck squamous carcinoma cells (HN31) and correlate transmembrane voltage with the cell volume. After discovering that cell volume is modulated by the membrane potential, we sought a less intrusive manner by which we could modify the cell’s electrical environment. For example, changing the concentration of an ionic species in a cell’s environment may change the cell’s membrane potential [25, 26]. In this case, since the membrane potential is not enforced by means of the patch clamp technique, the cell is now able to modify its internal ionic content and re-adjust its membrane potential. We can thus measure the volume of suspended cells and to determine how cell size is affected by changes in the ionic environment. We also use a microfluidic compression device [27] to hold a non-adherent cell in place, and measure cell volume in parallel with changes in the cell environment. We also investigate the role of the actin cytoskeleton in volume regulation. In parallel, we develop a mathematical model to explain cell volume as a function of transmembrane voltage and ionic content. Active ion pumps as well as passive channels and cotransporters are involved in ionic fluxes across the membrane. We propose from both experimental data and the model that the cell volume is mainly regulated by the total amount of intracellular ions and proteins; ion contents can be perturbed by the membrane potential or extracellular medium. As the protein content grows, the cell volume scales proportionally.

**Figure 1:**
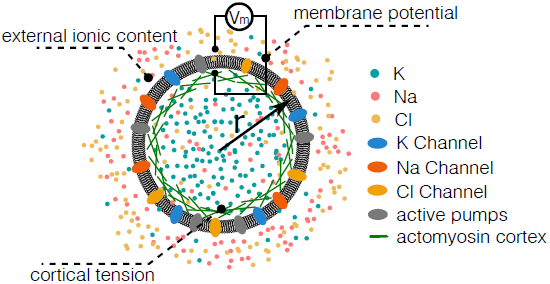
Cell volume regulation on short time scales is determined by ionic regulation. Major ion species in the cell are Na^+^, K^+^, and Cl^−^, which are all present in millimolar concentrations. These ion concentrations are controlled by passive ion channels and active ion pumps. Changes in ionic content also influence the transmembrane potential. In experiments, the membrane potential can be controlled by introducing external currents using a voltage clamp. The cell volume, which is determined by the overall water and protein content, can be modulated by changes in the transmembrane potential, external ionic content, or cortical tension.

## 2 Materials and Methods

### 2.1 Cell culture

HN-31, a head and neck squamous cell carcinoma (HNSCC) metastatic cell line was cultured in DMEM (Thermo Fisher Scientific, 11960044) supplemented with 1% L-glutamine (Lonza Group Ltd., Basel, Switzerland 17-605E), 0.5% penicillin-streptomycin solution (HyClone, thermo scientific SV30010), 2% MEM vitamin solution (Thermo Scientific - SH30599.01), 1% sodium pyruvate solution (Hyclone SH30239.01), 1% non-essential amino acids (Lonza Group Ltd., Basel, Switzerland13-114E), and 10% FBS. Cells were incubated at 37°C with 5% CO_2_ atmosphere and grown to ∼70-80% confluence.

**Table 1:** Experimental solutions (mM)

|  | Internal<br>patch | Modified<br>Ringer's | Low<br>Cl<sup>-</sup> | High<br>K<sup>+</sup> | Actin<br>depoly. |
| --- | --- | --- | --- | --- | --- |
| NaCl | 0 | 140 | 0 | 2.8 | 140 |
| Na-aspartate | 0 | 0 | 140 | 0 | 0 |
| K-glutamate | 135 | 0 | 0 | 0 | 0 |
| KCl | 0 | 2.8 | 2.8 | 140 | 2.8 |
| MgCl<sub>2</sub> | 5 | 1 | 1 | 1 | 1 |
| CaCl<sub>2</sub> | 0 | 2 | 2 | 2 | 2 |
| HEPES | 10 | 10 | 10 | 10 | 10 |
| NaOH | 0 | 1.5 | 1.5 | 1.5 | 1.5 |
| KOH | 9.4 | 0 | 0 | 0 | 0 |
| Na-ATP | 5 | 0 | 0 | 0 | 0 |
| EGTA | 10 | 0 | 0 | 0 | 0 |
| Lat. A | 0 | 0 | 0 | 0 | 10<sup>-3</sup> |

### 2.2 Experimental solutions

The bath solution used during voltage clamp by whole-cell patch clamp recording was a modified Ringer’s solution chosen to resemble the physiological environment. It consisted of (in mM): 140 NaCl, 2.8 KCl, 1 MgCl_2_, 2 CaCl_2_, and 10 HEPES. The pH was titrated to 7.25 with NaOH (final concentration ∼ 1.5 mM), and the osmolality was adjusted to 300 mOsm /kg by addition of glucose (15 mM). The internal solution used to fill the quartz pipettes was chosen to mimic the physiological contents of the cell. It consisted of (in mM) 135 K-glutamate, 5 MgCl_2_, 5 Na_2_-ATP, 10 EGTA, and 10 HEPES. The pH was adjusted to 7.25 with KOH (final concentration 9.4 mM), and the osmolality was 316 mOsm/kg. The potassium glutamate was made by titrating glutamic acid with potassium hydroxide. All chemicals were purchased from Sigma Aldrich.

Microfluidic solutions were based on the modified Ringer’s solution, with additional experiment-specific modifications: The low chloride solution was identical to the modified Ringer’s solution, except the 140 mM NaCl were substituted with 140 mM Na-aspartate, since aspartate is a negatively charged amino acid, too large to pass through most ion channels and pumps. For the high potassium solution, the amounts of KCl and NaCl were switched, so that the solution consisted of 2.8 mM NaCl and 140 mM KCl. For the actin depolymerization experiments, 1 μM Latrunculin A (Lat. A) was added to the modified Ringer’s solution. A detailed list of the solution content is in Tab. 1.

### 2.3 Microscopy

For voltage-clamp whole-cell patch clamp experiments, HN-31 cells were imaged with bright-field illumination (oil immersion 100x objective lens on a Axiovert 135 microscope; Zeiss, Oberkochen, Germany). A video camera (model No. NC-70x; DAGE-MTI, Michigan City, IN) and videocassette recorder (model No. AG-1970, S-VHS; Panasonic, Secaucus, NJ) were used to monitor and record (33 frames/second) the transmission image of cells over the duration of the experiment. The images were digitized and used to obtain the morphology of the cell (i.e., cell radius). For each frame of each experiment, a circle, parameterized by its radius and center, was fit to the cell by hand. Cells whose images could not be parameterized as a circle (i.e. cells with bleb formations) were excluded from the analysis.

For microfluidic experiments, HN31 cells were imaged with a confocal microscope (oil immersion 63 objective LSM 780 microscope; Zeiss). Cells were illuminated with a 405 nm photo-diode, and the emitted 460 nm and 580 nm fluorescence were detected by photomultiplier tubes. Images were acquired every 4 minutes to avoid photobleaching during the course of the experiment. After obtaining 80 minutes of time series images, confocal slices 1 micron apart were obtained for every cell.

For the chloride experiment on suspended cells, a 60x microscope (Nikon) and camera (Aven 26100-230 miniVUE Camera) were used to image the suspended cell every 15 s.

### 2.4 Whole-cell voltage-clamp experiments

For each voltage-clamp experiment the cells were trypsinized, centrifuged and re-suspended in fresh media. The cells were re-seeded on top of a 35 mm poly-D-lysine-coated glass bottom Petri dish (MaTek Corp., Ashland, MA) suspended in modified Ringer’s solution. Experiments were conducted before the cells attached to the substrate, within about 10 mins after re-seeding. This ensured that the cells remained spherical, allowing estimation of volume from radius measurements. The Petri dish was perfused with Ringer’s solution with a peristaltic pump (model No. RP-1 4 channel pump; Rainin, Oakland, CA) at a constant flow rate of 0.5 mL/min.

Quartz pipettes were pulled with a laser-based puller (Sutter Instrument, Novato, CA), coated with Sylgard (Dow Corning, Midland, MI) and had resistances of less than 10 MΩ. A single suspended cell was chosen for an experiment. The suspended cell was attached to the quartz pipette by applying negative pressure to the pipette, and the pipette pressure was controlled during giga-seal formation as well as during recordings with a high-speed pressure clamp (HSPC-1; ALA Scientific Instruments, Farming-dale, NY). Voltage clamp of the cell’s membrane potential was maintained with an amplifier (Axon 200B, Molecular Devices, Union City, CA) configured in whole-cell patch-clamp mode. Voltages were measured relative to a reference electrode (Ag/AgCl) that was placed in the bath. The holding potential was maintained while recording the cell morphology, where the pipette pressure was held approximately constant at −0.9 mmHg. For the entire duration of the recording, a 5 mV amplitude square-wave was added to the holding potential to monitor the DC conductance of the cells. The voltage stimulus and data acquisition were controlled by software written in Lab-VIEW for Windows (v8.5.1) in conjunction with a digital acquisition card (PCI-6052E, National Instruments, Austin, TX), and the data analysis were preformed using MATLAB (The Math-Works, Natick, MA).

### 2.5 Microfabrication of microfluidic compression device

The compression device is able to maintain the cells in the same position under external fluid flow [27]. Silicon wafers were patterned with SU8-2100 (MicroChem), using UV exposure and photomasks (Fineline Imaging, Inc., CO). Wafers were created for the cell and air chambers, which were fabricated with PDMS (Sylgard 184, Dow Corning Corp.). Using another mask and UV exposure, micropillars 10–15 microns tall were printed directly onto a glass coverslip from SPR220-7, using the adhesion promoter Hexamethyldisilazane (HMDS, Sigma-Aldrich). Holes were punched into the air chamber layer, which was then aligned with the cell chamber layer. The layers were bonded by baking at 80°C. The inlet and outlet holes to the cell chamber were then punched into the PDMS, the cell and air chambers were aligned with the micropillar layer, and the PDMS was bonded to the coverglass with plasma treatment and baking at 80°C overnight before use.

### 2.6 Microfluidic experiments

Microfluidic devices were incubated with 100 mg/mL BSA prior to cell loading in order to decrease cell adhesion to the glass. After remaining in the microfluidic device for at least 30 minutes, the BSA was removed from the device, and the cell chamber was rinsed with modified Ringer’s solution.

The microfluidic device was arranged on the microscope stage, and the air pressure in the device’s air chamber was adjusted to 25 psi in order to compress the cells. An electric syringe pump (Harvard Apparatus, MA) was attached to the cell chamber inlet and outlet in order to induce solution flow through the cell chamber. After compression, the flow was set to ∼80 *μL*/minutes to flush out any small, uncompressed cells and cell debris and to continuously refresh the cells’ environment.

After about 30 minutes, the external solution supplied by the syringe pump was changed to low-sodium/high-potassium solution.

### 2.7 Chloride concentration experiments

HN31 cells grown in culture were trypsinized and resuspended in fresh media. After centrifugation, cells were resuspended in the modified Ringer’s solution described above. Glass, fire-polished pipettes were filled with Ringer’s solution and were attached to micromanipulators. The suspended cells were attached to the pipette by applying negative pressure inside the pipette. Unlike the patch-clamp experiment, the chloride concentration experiments did not use the whole-cell voltage clamp configuration; rather, the pipette was simply used to hold a single cell in suspension while measuring its size. During this process, the cell bath was slowly refreshed with the low chloride solution at a flow rate of ∼3 μL/minutes.

## 3 Mathematical Model

Here we present a mathematical model that relates cell’s electrical physiology to the cell volume through the regulation of ion fluxes and intracellular osmotic content. This model is motivated by the observation that the cell volume is correlated to the transmembrane voltage and extracellular ionic environment. The model focuses on osmolarity regulation through ionic contributions on short time scales, and ignores longer time scale changes in cell cytoskeletal organization and contractility. By their relative contributions to the intracellular osmolarity, we consider four ionic species, Na^+^, K^+^, Cl^−^, and A^−^, in the model, where A^−^ represents negatively-charged large molecules or proteins that are not permeable across the cell membrane. The extracellular ion concentrations 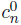 are given according to the specification of each experiment. Here *c* represents concentrations, the superscript ‘0’ indicates quantities associated with extracellular environment, and *n* ∈ {Na^+^, K^+^, Cl^−^, A^−^}. The total number of intracellular proteins *N*_*A*_ is given, but its effective valence is unknown and is thus assumed. Without loss of generality, we assume the average valence of t hesemolecules to be −1.

At short time scales (∼ minutes), changes in the cell size are mainly determined by water transport; therefore the cell radius, *r*, is given by [15]

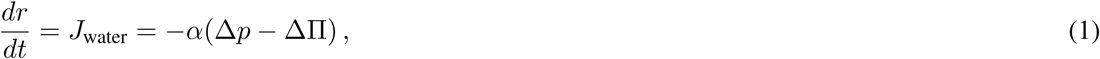

where *J*_water_ is water flux across the cell membrane, defined positive inward. *α* is the coefficient of water permeation. Δ? is the osmotic pressure difference across the cell surface. The hydrostatic pressure difference across the cell surface, Δ*p*, is related to the cortical stress, σ, by the Laplace law

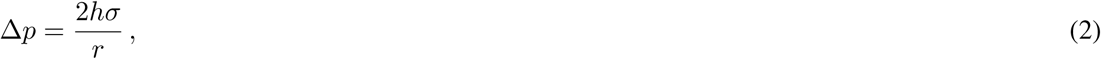

where *h* is the thickness of the cell cortex. The cortical stress can be solved from constitutive equations of the actomyosin cortex [15]

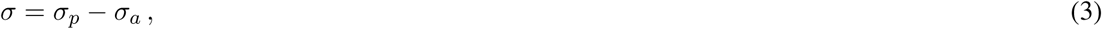

where σ_*p*_ is the passive mechanical stress of the actin network, which includes contributions from F-actin crosslinkers and filament mechanics, and σ_*a*_ is the active myosin contraction from the cortex. The passive stress is typically small when compared to the active stress, and in this model, we consider σ_*p*_ as a parameter.

The governing equations for the total amount of each intracellular ion species, *N*_*n*_, is

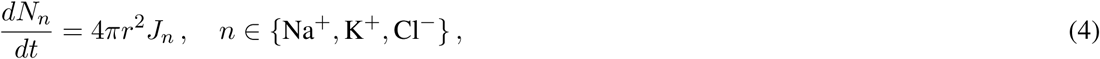

where *J*_*n*_ is the total ion flux across the membrane for each species and is determined by the boundary conditions of ion fluxes through the membrane channels. In general, ions are transported across membranes both passively and actively. Passive ionic transport is carried by ion-specific channels and pores, some of which are gated by membrane tension, *τ_*m*_* = *σh*. Active transport is carried by energy-consuming ion pumps, which utilize chemical energy (ATP) to transport ions against a chemical potential gradient. Both modes of transport are incorporated in our model:

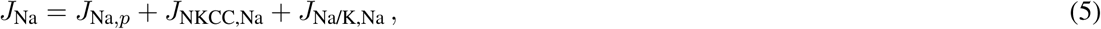

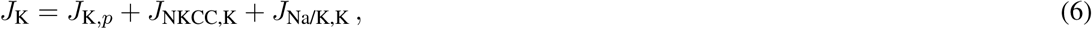

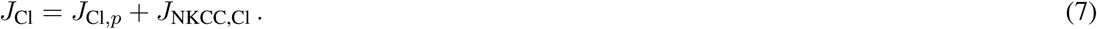

The passive ion fluxes, *J*_*n,p*_, are proportional to the electrochemical potential difference of ions across the membrane [28],

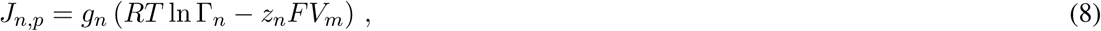

where 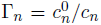is the ratio of extra- and intra-cellular ion concentrations. *g*_*n*_ = *G*_0,*n*_ *T*_*m*_ is the rate of ion permeation of each species, where *G*_0,*n*_ is a constant depending on the channel property and the density of the channels. *T*_*m*_ ∈ (0, 1) is a mechanosensitive gating function that follows a Boltzmann distribution, i.e.,

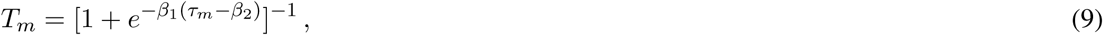

where *β*_1_ and *β*_2_ are two constants.

Another passive channel to consider is the Na^+^-K^+^-Cl^−^ cotransporter (NKCC) which, along with its isoforms, is widely expressed in various cell types [29]. The NKCC simultaneously transports one Na^+^, one K^+^, and two Cl^−^s into the cell under physiological conditions, the flux of which can be written as [30]

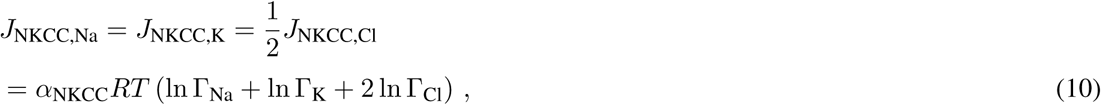

where *α*_NKCC_ is a transport rate constant independent of the membrane tension. Since transported Na-K-Cl_2_ is electrically neutral, its flux is independent of the membrane potential.

For the active fluxes, we consider the Na^+^/K^+^ pump, a ubiquitous and important ion pump in animal cells. It exports three Na^+^ ions and intakes two K^+^ ions per ATP unit. Because the overall flux is positively charged, the activity of the pump depends on the membrane potential [31]. In addition, the flux depends on the concentrations of Na^+^ and K^+^ [32, 33] and saturates at high concentration limits [33]. By decoupling the dependence of the voltage and ion concentration, as a modification of existing models [33, 34], we express the flux of Na^+^ and K^+^ through the Na^+^/K^+^ pump as

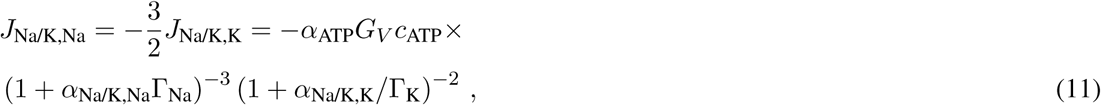

where *α*_ATP_ is a transport rate constant of the pump, *c*_ATP_ is the concentration of ATP. *α*_Na/K,Na_ and *α*_Na/K,K_ are constants that scale Γ_Na_ and Γ_K_, respectively. The exponents 3 and 2 are the Hill’s coefficients of Na^+^ and K^+^, respectively. Equation 11 ensures that the flux is zero when either 1*/*Γ_Na_ or Γ_K_ goes to zero; the flux saturates if 1*/*Γ_Na_ and Γ_K_ go to infinity. *G*_*V*_ captures the voltage-dependence of the pump activity [31]

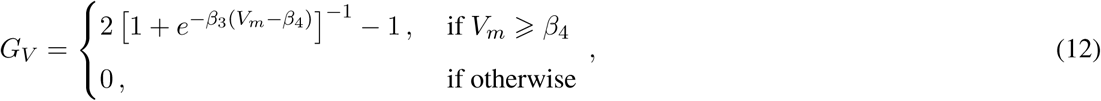

where *β*_3_ and *β*_4_ are constant.

The equation for the membrane potential is solved by the electro-neutral condition, i.e.,

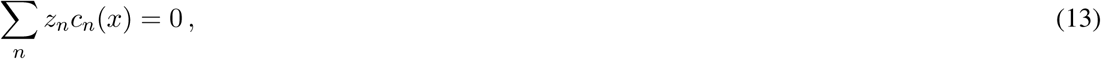

where *z*_*n*_ is the valence of each ionic species. The electro-neutral condition also applies to extracellular medium. In addition to the experiment-specified 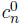, we also add either positively or negatively charged large molecules to balance the total charge in the medium. This added molecules contributes to extracellular osmolarity as well.

For the steady-state solution, *d/dt* = 0. In the general problem, the unknowns are *r, c*_Na_, *c*_K_, *c*_Cl_, *V*_*m*_. Five equations 1, 4, and 13 are used to solve the five unknowns.

Under a voltage clamp, the unknowns of the system are *r, c*_Na_, *c*_K_, *c*_Cl_, *J*_clamp_ and Eqs. 5-7 are replaced

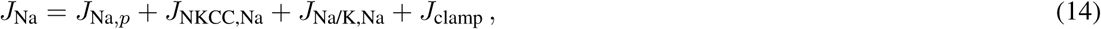

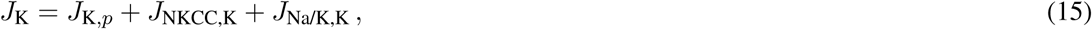

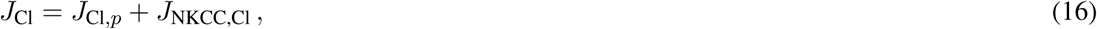

where *J*_clamp_ is an additional current introduced by the voltage clamp. Note that this current is different from adjusted ion flux balance under voltage clamp.

Unless otherwise specified, the parameters in the model are listed in Tab. 2.

## 4 Results

### 4.1 Cell volume increases with direct depolarization

We used whole cell patch clamp to obtain electrical access to the cytoplasm of the cell. Using suspended cells enabled us to measure cell size with bright field illumination, from which cell volume could be estimated due to their spherical geometry. For the initial experiments the membrane potential of the cell was clamped at −60 mV, a typical resting potential for cells [35], for about 20 minutes, allowing the volume of the cell to equilibrate. We then changed the potential to either −120 mV, −90 mV, −30 mV, or 0 mV. After every change in potential, we waited 10–20 minutes for the cell volume to readjust. A typical cell completed its response to voltage change in 10 minutes (Fig. 2A). We find that the cell volume increases monotonically with increasing transmembrane voltage (Fig. 2B). The cell overall volume change is roughly 50% when the membrane potential is zero (Fig. 2B).

**Table 2:** Membrane channel related parameters used in the electromotility model.

| Parameters | Description | values |
| --- | --- | --- |
| $r_0$ ($\mu\text{m}$) | Initial cell radius | 9.1 |
| $h$ (nm) | Cortical thickness | 10 |
| $\sigma_p$ (Pa) | Passive membrane stress | $1 \times 10^2$ |
| $\sigma_a$ (Pa) | Active contraction | $-3 \times 10^3$ |
| $D_{\text{Na}}$ ($\text{m}^2/\text{s}$) | Diffusion constant of Na | $1.33 \times 10^{-9}$ |
| $D_{\text{K}}$ ($\text{m}^2/\text{s}$) | Diffusion constant of K | $1.96 \times 10^{-9}$ |
| $D_{\text{Cl}}$ ($\text{m}^2/\text{s}$) | Diffusion constant of Cl | $2.03 \times 10^{-9}$ |
| $D_{\text{A}}$ ($\text{m}^2/\text{s}$) | Diffusion constant of A | $5.0 \times 10^{-10}$ |
| $N^-$ (mol) | Total intracellular A | $3.9 \times 10^{-13}$ |
| $\alpha$ (m/s/Pa) | Water permeability | $3.0 \times 10^{-11}$ |
| $G_{0,\text{Na}}$ ($\text{mol}^2/\text{Kg}/\text{m}^3$) | Na channel constant | $8 \times 10^{-12}$ |
| $G_{0,\text{K}}$ ($\text{mol}^2/\text{Kg}/\text{m}^3$) | K channel constant | $1.6 \times 10^{-9}$ |
| $G_{0,\text{Cl}}$ ($\text{mol}^2/\text{Kg}/\text{m}^3$) | Cl channel constant | $4 \times 10^{-10}$ |
| $c_{\text{ATP}}$ ($\text{mol}/\text{m}^3$) | ATP concentration | 1 |
| $\alpha_{\text{ATP}}$ (m/s) | Coefficient in $J_{\text{Na/K}}$ | 0.1 |
| $\alpha_{\text{Na/K,Na}}$ | Constant in $J_{\text{Na/K}}$ | 0.01 |
| $\alpha_{\text{Na/K,K}}$ | Constant in $J_{\text{Na/K}}$ | 0.1 |
| $\alpha_{\text{NKCC}}$ ($\text{mol}/\text{m}^2/\text{s}$) | Coefficient in $J_{\text{NKCC}}$ | $1 \times 10^{-10}$ |
| $\beta_1$ | Constant in $T_m$ | 10 |
| $\beta_2$ (N/m) | Constant in $T_m$ | $3.0 \times 10^{-5}$ |
| $\beta_3$ | Constant in $G_V$ | 1.5 |
| $\beta_4$ (mV) | Constant in $G_V$ | -92 |

**Figure 2:**
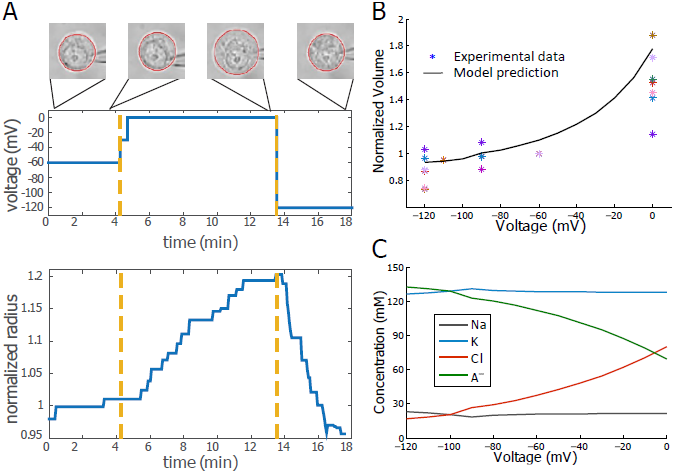
Cell volume correlates with transmembrane voltage. (A) As the membrane holding potential is changed (upper panel), the cell radius and volume changed with time (lower panel). The data is shown for one cell for illustration. (B) The measured cell volume vs. transmembrane voltage under steady-state. Cell volume is normalized with respect to its volume at −60 mV. The volume increases with depolarization. Star: experimental data. Different colors represent different cells (*N* = 10). Solid line: model prediction. (C) Model perdition on the intracellular ionic content of the four species as functions of membrane potential.

The observed relation between membrane potential and cell volume can be predicted by our electromechanical model (Fig. 2B). The model suggests that as the cell is depolarized and volume increases, the concentrations of Na^+^ and K^+^ almost remain the same (Fig. 2C), showing that in additional to Cl^−^ influx upon depolarization, Na^+^ and K^+^ also flow into the cell. An increased concentration of Cl^−^ suggests a decreased concentration of A-since the electro-neutral condition must be satisfied. The decreased concentration of A^−^can only happen when the cell volume increases, as seen in the experiment and predicted by the model.

### 4.2 Cell volume decreases with decreased extracellular chloride concentration

Since ion flux is involved in cell volume regulation, we next ask if it is possible to control cell volume by changing the chemical potential of membrane-permeable ions in the cell’s environment. Given that chloride is the only abundant negatively charged ions in the cell medium, we would like to examine the effect of extracellular chloride concentration on the cell volume. In this experiment we replaced 140 mM of chloride in the modified Ringer’s solution with aspartate, a negatively charged ion that is too large to pass through the cell membrane (Fig. 3A). In order to avoid effects of shear stress from the fluid onto the cell, we gradually replaced the high concentration chloride solution with the high concentration aspartate solution instead of implementing a step change in solution.

**Figure 3:**
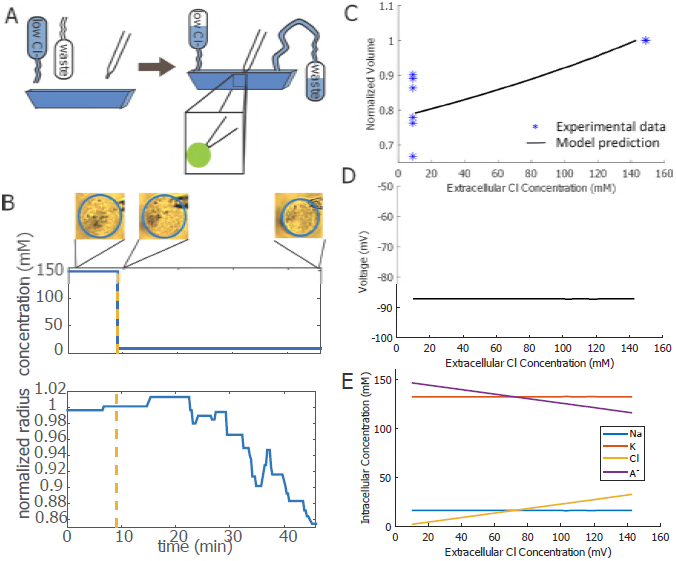
Cell volume correlates with extracellular chloride concentration. (A) Cells are suspended using a micropipet, the extracellular medium is changed to low chloride solution, and the cell volume is simultaneously monitored. The pipette was simply used to hold a single cell in suspension while measuring its size. (B) The measured cell volume for a single HN31 cell as a function of time during chloride solution switch. The upper panel indicates the time of medium switch; since the low chloride medium gradually flew into the solution, the exact profile of chloride concentration is not a perfect step function. (C) Cell volume shows a ∼10% decrease as chloride is decreased in the extracellular media. Star: experimental data, *n* = 6, MEAN ± Std. = 0.81 ± 0.09. Solid line: model prediction. (D) Model prediction of membrane potential as a function of extracellular chloride concentration. (E) Model perdition on the intracellular ionic content of the four species as functions of membrane potential. In this model, *α*_NKCC_ = 1.5 × 10^−10^ mol/m^2^/s to account for the cellular response to medium shock with low chloride.

Fig. 3B shows the response of a typical cell to low-chloride medium. About 30 minutes after switching to the aspartate solution, the cell radius had decreased by about 10%. The time scale for cell volume adjustment during the chloride/aspartate experiment was roughly three times longer than the time scale for cell volume adjustment during the voltage clamp experiment. This is likely due to the fact that the bath solution changed from the high-chloride solution to high-aspartate solution over a period of about fifteen minutes, whereas the membrane potential changed almost instantaneously during the voltage clamp experiment. Figure 3C shows the steady-state volume of cells that were switched into a low chloride solution after 30 minutes. Lowering the chloride concentration in the cell environment led to cell volume shrinkage of 5–10%.

This volume reduction can be predicted by our model (Fig. 3C). Unlike the voltage clamp case where the membrane potential and cell volume is correlated (Fig. 2B), here with the medium chloride depletion the membrane potential is not predicted to vary (Fig. 3D). The intracellular chloride concentration is predicted to decrease in low-chloride medium (Fig. 3E). This is because the chemical potential of chloride in the medium decreases so that chloride is more likely to flux out of the cell. In addition, the model predicts constant intracellular sodium and potassium concentrations when external chloride concentration reduces, indicating efflux of both in the process since the cell volume is decreased.

### 4.3 Cell volume increases with sodium and potassium exchanged in the medium

In addition to chloride, the cell and its medium are rich in sodium and potassium. Under physiological conditions the ratio of intracellular to extracellular concentrations of sodium is less than one-tenth while that of potassium is more than ten. A perturbation of these ratios may have an effect on cell volume. We therefore examined such effect and exchanged the extracellular content of sodium and potassium by slowly flowing low-sodium/high-potassium medium into a microfluidic device to replace the original Ringer’s solution (Fig. 4A). A slight compression from the devices kept the cells in place without over-deforming the spherical cell shape (Fig. 4A).

**Figure 4:**
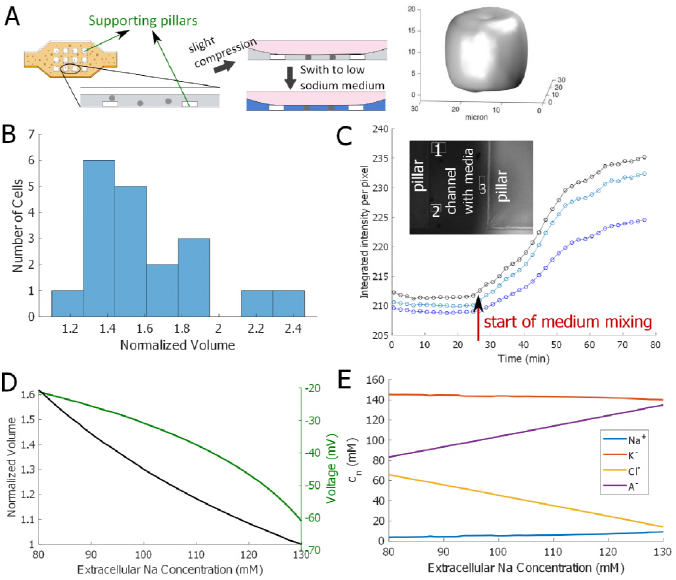
(A) Cells are slightly compressed in the compression microfluidic device and held in place to avoid being washed away by the medium. The supporting pillars were used to support the compression chamber and to avoid cells being over compressed. A confocal Z-stack reconstruction of the cell demonstrated that the cells were still close to spheres. (B) Experimental data showing that cell volume increased in the low-sodium/high-potassium medium. *n* = 19, MEAN ± Std. = 1.58 ± 0.33. (C) New media with dye was flown into the device, and different regions of the device were imaged. The dye time series data for different regions of the device shows that mixing is not uniform. Here we show the dye intensity at three sample nearby locations in the channel. (D) Model prediction of cell volume and membrane potential as the external sodium is gradually replaced by potassium. (D) Model prediction of intracellular ionic contents as the external sodium is gradually replaced by potassium. In this model, the following parameters are adjusted to account for the cellular response to medium shock. *α*_ATP_ = 1 m/s, *α*_Na/K,Na_ = 5, *α*_Na/K,K_ = 0.01, *α*_NKCC_ = 1 × 10^−13^ mol/m^2^/s. These parameters are mostly involved in the Na/K pump.

On average the cell volume increased by about ∼60% in the low-sodium/high-potassium medium (Fig. 4B). Due to the non-uniform mixing rate in the microfluidic device (Fig. 4C), it is possible that not all cells experienced the solution exchange at the same time and to the same degree. It is therefore reasonable to expect that there is a wide variation in cell response in volume.

This volume increase can again be predicted by our model (Fig. 4D). Although a depletion of chloride in the medium does not lead to a predicted membrane potential change (Fig. 3D), our model does predict cell depolarization when cells are exposed to low-sodium/high-potassium medium (Fig. 4D). This result is consistent with experimental findings that cells become depolarized when surrounded by a high potassium environment [36]. Our model predicts that the intracellular concentrations of sodium and potassium follow slightly with the trend of the corresponding extracellular content (Fig. 4E). This can be interpreted from the change of the extracellular chemical potential of the two ionic species. Similar to the model prediction for the voltage clamp (Fig. 2C), here we also see a predicted large chloride influx upon the cell depolarization (Fig. 4E).

### 4.4 Cell volume increases with actin depolymerization

In order to decouple the effects of cytoskeletal forces and osmotic forces on maintaining cell volume, we used 1 μM Lat. A to disrupt the actin cytoskeleton (Fig. 5A). Once again, we used the compression device to hold cells in place while the surrounding medium was replaced, this time with the Lat. A solution. On average, the cell volume increased by ∼6% upon actin depolymerization (Fig. 5B). Similarly, because of the non-uniform mixing rate in the microfluidic device, we see a wide spread of volume change across cells.

**Figure 5:**
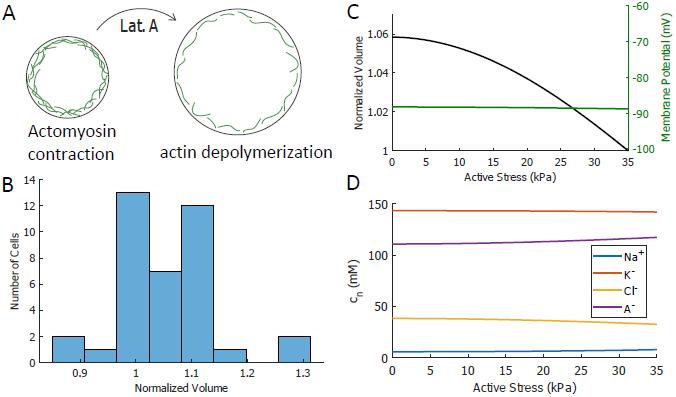
Role of actin cortex in maintaining cell volume. (A) Schematics showing depolymerizing actin disrupts the cortex and reduces active contractile stress in the cortex, leading to cell volume increases (not to scale). (B) Experimental data showing that cell volume increases upon actin depolymerization. *n* = 38, MEAN ± Std. = 1.06 ± 0.09. (C) Model prediction on cell volume and membrane potential when Lat. A is added to the cell. (D) Model prediction of the concentrations of intracellular species. *α*_ATP_ = 0.25. In this model we let the effective membrane thickness vary linearly from 500 nm to 10 nm as the contraction stress decreases.

In the model, the effect of Lat. A treatment can be reflected in a reduction of the active cortical stress, σ_*a*_. In addition, since actin cortex contributes to the effective membrane thickness, actin depolymerization reduces the thickness of this layer as well. In combination, these lead to a reduced cortical tension, σ*h*, in the Laplace equation (Eq. 2). Figure 5C shows the model prediction of cell volume increase under actin depolymerization. Since the cortical tension decreases while cell volume increases, the right hand side of Eq. 2 decreases, meaning that the hydrostatic pressure difference across the cell reduces. Under steadystate where the chemical potential of water is balanced, a reduced intracellular hydrostatic pressure means a reduced intracellular osmotic pressure. We will see in the next section that the relative magnitude of this change is very small and the intracellular pressure is almost a constant.

While volume increases, the membrane potential is predicted constant (Fig. 5C) since actin depolymerization probably does not interfere with the electrical potential. The intracellular ionic variation (Fig. 5D) is a combined effect of electro-neutral condition and water chemical potential balance across the membrane.

### 4.5 Cell volume increases with intracellular ion and protein content

Finally, we present model predictions on cell volume with varying intracellular content. We are particularly interested in the cytoplasmic protein content (A^−^ in the model) because as cells grow and age, the protein content increases as well. Our model predicts that the cell volume increases almost linearly with the total amount of cell proteins (Fig. 6A); however, the concentration of the protein, as well as other ion species, remains constant (Fig. 6B). We can see that the extensive properties of the cell, such as cell volume and protein content, scale linearly with each other, while intensive properties, such as ion and protein concentrations and membrane potential (Fig. 6A), remain the same. The cell maintains almost constant osmolarity and hydrostatic pressure as it grows for a fixed active cortical tension.

**Figure 6:**
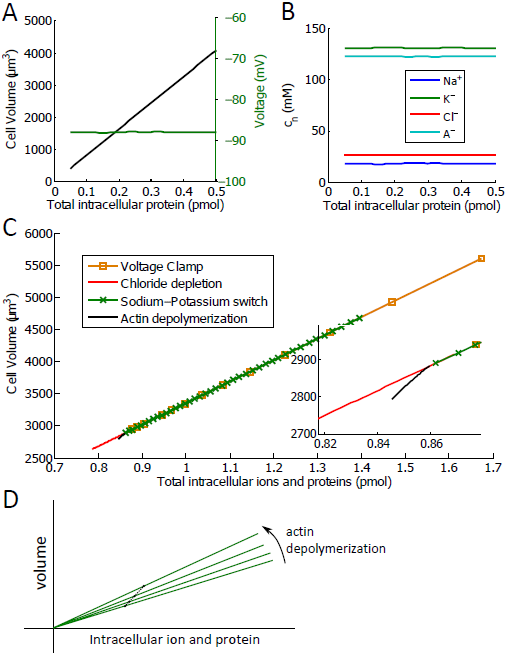
(A) Model prediction of the cell volume (left axis) and voltage (right axis) as a function of intracellular protein content. (B) Model prediction of the intracellular ion and protein concentration with intracellular protein content. (C) Model prediction of the cell volume with the combined total intracellular ion and protein content from the four experiments: voltage clamp, low-chloride medium, low-sodium/high potassium medium, and actin depolymerization. Inset: zoom-in for better visualization of the curve for actin depolymerization. (D) Schematic explaining the volume-content relation during actin depolymerization (not to scale). Each solid curve represents a generic volume-content relation for a fixed cortical tension, similar to the chloride depletion curve, for example, in panel C. A trajectory connecting different points on different solid lines gives the actin depolymerization curve in panel C.

We can check this idea by examining how cell volume scales with the intracellular ion and protein contents in three situations examined experimentally: voltage clamp, low-chloride medium, and low-sodium/high potassium medium. Fig. 6C shows that the model predicts that the cell volume vs. total cytoplasmic content for these three cases fall on the same line, indicating that the cell volume scales with the total intracellular ion and protein contents. For the case with Lat. A treatment, the cell volume still scales with the total intracellular ion and protein contents but the scaling factor, i.e., the slope, changes slightly. This change can be seen after zoom-in but the magnitude of the change is almost negligible.

The exact slopes in Fig. 6C can be estimated from the model. Let *V* = 4*πr*^3^*/*3 be the volume of the cell and *M* = *V* ∑_*n*_ *c*_*n*_ be the total intracellular ion and protein content. By Eq. 1 under steady-state,

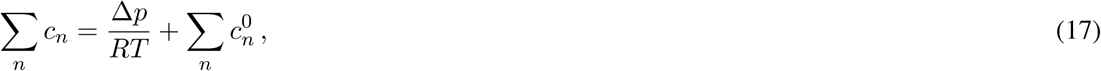

where *n* ∈ {Na^+^, K^+^, Cl^−^, A^−^}. Substituting Eq. 2 into the above equation, we have

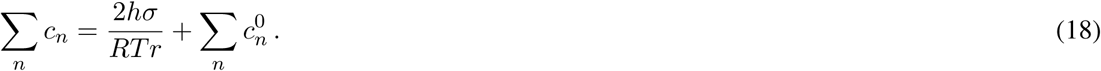

This relationship implies that the total intracellular concentration is only determined by the extracellular concentration, cortical tension, and cell volume. If the cortical tension scales with the volume of the cell, then the internal concentration is a constant, independent of volume. The slope of the volume-content relation can be calculated as

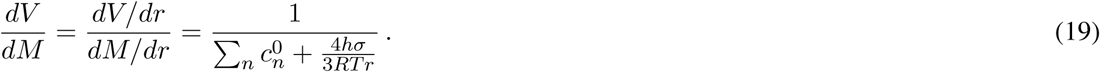

Using order of magnitude estimates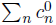 is on the order of 300 mM and 4*hσ*/3*RTr* may vary from 10^−3^ mM to 10 mM, much smaller than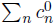. Therefore, the slope is roughly constant for fixed external total ionic concentration, leading to the almost linear relation in Fig. 6C. Upon actin depolymerization, the slope will increase slightly as illustrated in Fig. 6D. A trajectory connecting different points of different cortical tensions gives the actin depolymerization curve in Fig. 6C.

## 5 Discussions and Conclusions

In this paper, using patch clamp experiments and microfluidic devices we explored how individual HN31 cell volume is correlated with the ionic environment of the cell, the transmembrane voltage, and the active stress from the actin cortex. While the exact results may vary from cell line to cell line, here we focus on a generic response of cells under these conditions. An electrophysiology model is presented to facilitate the generic understanding of cell volume change under these experimental conditions.

We found that changes in the transmembrane electrical potential and ionic environment of the cell can influence cell volume. For example, in the voltage clamp experiment we see a direct correlation between cell volume and membrane potential—depolarized cells have larger volumes, which was also seen in barnacle muscle cells [37]. In the low-sodium/high-potassium medium the membrane potential is predicted to change by at least 50%. This membrane potential change does not always happen in other conditions. For example, in the external chloride depletion and actin depolymerization experiments, the membrane potential is predicted to be unchanged. The relationship between membrane potential and cell volume is mediated by the ion channels and pumps, i.e., the membrane potential change leads to a significant ion fluxes and the cell adjusts its volume by maintaining the same intracellular osmolarity. It is important to note that for all known ion channels and pumps, the flux only depends on the intracellular and extracellular ion concentrations and the transmembrane voltage (which itself is a function of ion concentrations), and not a function of the total ionic content. This property leads to homeostatic ion concentration differences at steady state and ultimately determines how total ionic and protein content scales with cell volume.

Voltage clamp is not the only way to perturb ion fluxes—a change of external ionic environment can also accomplish the same. In this work, we have seen that modified external ionic environment can change cell volume through changing ion fluxes but does not always affect the membrane potential. The exact volume change depends on ion fluxes and thus on ion pump activities. It is possible that for different cell types the volume change can be quantitatively or even qualitatively different. In this model the cell maintains almost constant intra-to extracellular sodium and potassium concentrations due to the active Na^+^/K^+^ pump. When this pump is disabled, the two concentrations can vary significantly.

The model predicts significant change of membrane potential when cells are in the low-sodium/high-potassium medium (Fig. 4D), in contrast to the constant membrane potential in low-chloride medium (Fig. 3D). This difference comes from the predicted non-trivial change of intra-to extracellular chloride concentration ratio in the low-sodium/high-potassium medium (Fig. 4E). Under steady-state the total ion flux of each species is zero, i.e., Eqs. 5–7 vanish. We can re-arrange the three equations to eliminate *J*_Na/K_ and *J*_NKCC_, obtaining

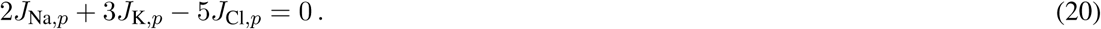

Substituting Eq. 8 into Eq. 20, we obtain an expression for the membrane potential

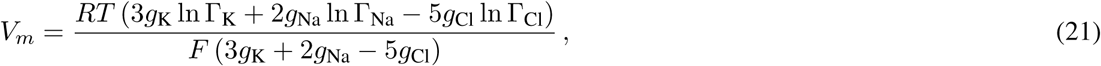

which can be regarded as a revised Goldman-Hodgkin-Kate equation [25] when Na^+^/K^+^ pump and NKCC are considered. In the low-chloride medium case, Γ_K_ and Γ_Na_ remain the same (Fig. 3E) and Γ_Cl_ is also close to a constant since the intracellular chloride concentration decreases with extracellular chloride concentration (Fig. 3E). The membrane potential in Eq. 21 therefore remains constant. In the low-sodium/high-potassium medium case, the intracellular sodium and potassium concentrations vary with external concentrations (Fig. 4E) so that Γ_K_ and Γ_Na_ do not change significantly. However, the intracellular concentration of chloride increases by several folds upon medium switch (Fig. 4E), leading to a large decrease in Γ_Cl_, which in turn depolarizes the cell, as seen in Fig. 4D. This effect is mainly due to the presence of NKCC, and this prediction can be checked in experiments with sufficiently precise single cell voltage measurements.

Although active channel pumps regulate the intracellular ionic concentrations of each species and sometimes the membrane potential as well, the model actually predicts that they have no influence on the cell volume overall. For example, in the model we can remove the Na^+^/K^+^ pump and the NKCC and still retain the relation seen in Fig. 6C. This does not mean that active pumps do not play a role; rather, their role is mainly to regulate each *c*_*n*_ in a way that Σ_*n*_ *c*_*n*_ does not change significantly (Eq. 18). This result is a re-statement of the volume-content relation (Fig. 6C) that the intracellular protein content is one of the leading order influencers of cell volume. Note that changes in *c*_*n*_ would change functions of biologically important proteins such as the ribosome [38], and therefore the ion pumps are important in maintaining the proper chemical environment in which the cell can function.

Actin dynamics and cortical contractile stress can also affect cell volume through multiple mechanisms such as vesicle transport and tension-activated channels [39, 40]. In this paper we found that the actin cortex, on average, had a small effect on HN31 cell volume. It is likely that the effect of actin cortical tension on cell volume depends on the cell type. In general, there would be some cross-talk between actomyosin dynamics, membrane tension, ion channel distribution, and activation of ion channels that further connect actin structure and cell volume. The full picture of cell volume regulation on short time scales still awaits clear experimental and theoretical elucidation.

## References

[1] M. R. Bennett, W. L. Pang, N. A. Ostroff, B. L. Baumgartner, S. Nayak, L. S. Tsimring, and J. Hasty. Metabolic gene regulation in a dynamically changing environment. Nature, 454(7208):1119–1122, 2008.

[2] L. Lopez-Maury, S. Marguerat, and J. Bahler. Tuning gene expression to changing environments: from rapid responses to evolutionary adaptation. Nat. Rev. Genet., 9(8):583–593, 2008.

[3] A. N. Brooks, S. Turkarslan, K. D. Beer, F. Yin Lo, and N. S. Baliga. Adaptation of cells to new environments. Wiley. Interdiscip. Rev. Syst. Biol. Med., 3(5):544–561, 2011.

[4] A. C. Lloyd. The regulation of cell size. Cell, 154(6):1194–1205, 2013.

[5] S. Warntges, H. J. Grone, G. Capasso, and F. Lang. Cell volume regulatory mechanisms in progression of renal disease. J. Nephrol., 14(5):319–326, 2001.

[6] Xiaolong Yang and Tian Xu. Molecular mechanism of size control in development and human diseases. Cell Res., 21(5):715–729, 2011.

[7] S. Gonzalez and C. Rallis. The tor signaling pathway in spatial and temporal control of cell size and growth. Front. Cell Dev. Biol., 5:61, 2017.

[8] F. Lang, M. Ritter, N. Gamper, S. Huber, S. Fillon, V. Tanneur, A. Lepple-Wienhues, I. Szabo, and E. Gulbins. Cell volume in the regulation of cell proliferation and apoptotic cell death.˙ Cell Physiol. Biochem., 10(5–6):417–428, 2000.

[9] M. Jakab and M. Ritter. Cell volume regulatory ion transport in the regulation of cell migration. In Mechanisms and Significance of Cell Volume Regulation, volume 152, pages 161–180.

[10] F. Lang. Mechanisms and significance of cell volume regulation. J. Am. Coll. Nutr., 26(sup5):613S–623S, 2007.

[11] J. Tao, Y. Li, D. K. Vig, and S. X. Sun. Cell mechanics: a dialogue. Rep. Prog. Phys., 80(3):036601, 2017.

[12] M. P. Stewart, J. Helenius, Y. Toyoda, S. P. Ramanathan, D. J. Muller, and A. A. Hyman. Hydrostatic pressure and the actomyosin cortex drive mitotic cell rounding. Nature, 469(7329):226–230, January 2011.

[13] S. F. Pedersen and E. K. Hoffmann. Possible interrelationship between changes in F-actin and myosin II, protein phosphorylation, and cell volume regulation in ehrlich ascites tumor cells. Exp. Cell Res., 277:57–73, 2002.

[14] J. Nicholson, F. Korber, and S. Lambert. Application of moderate hydrostatic pressure induces unit-cell changes in rhombohedral insulin. Acta. Crystallogr. D Biol. Crystallogr., 52(5):1012–1015, 1996.

[15] H. Jiang and S. X. Sun. Cellular pressure and volume regulation and implications for cell mechanics. Biophys. J., 105(3):609–619, 2013.

[16] D. J. Blackiston, K. A. McLaughlin, and M. Levin. Bioelectric controls of cell proliferation: ion channels, membrane voltage and the cell cycle. Cell cycle, 8(21):3527–3536, 2009.

[17] B. T. Chernet and M. Levin. Transmembrane voltage potential is an essential cellular parameter for the detection and control of tumor development in a xenopus model. Dis. Model. Mech., 6(3):595–607, 2013.

[18] T.H. Hui, K.W. Kwan, T.T. Chun Yip, H.W. Fong, K.C. Ngan, M. Yu, S. Yao, A.H. Wan Ngan and Y. Lin. Regulating the membrane transport activity and death of cells via electroosmotic manipulation. Biophys. J., 110:2769–2778, 2016.

[19] G. Salbreux, J.-F. Joanny, J. Prost, and P. Pullarkat. Shape oscillations of non-adhering fibroblast cells. Phys. Biol., 4(4):268, 2007.

[20] G. L. Hunter, J. M. Crawford, J. Z. Genkins, and D. P. Kiehart. Ion channels contribute to the regulation of cell sheet forces during drosophila dorsal closure. Development, 141(2):325–334, 2014.

[21] E. K. Hoffmann, N. B. Holm, and I. H. Lambert. Functions of volume-sensitive and calcium-activated chloride channels. IUBMB Life, 66(4):257–267, 2014.

[22] Z. Qiu, A. E. Dubin, J. Mathur, B. Tu, K. Reddy, L. J. Miraglia, J. Reinhardt, A. P. Orth, and A. Patapoutian. SWELL1, a plasma membrane protein, is an essential component of volume-regulated anion channel. Cell, 157(2):447–458, 2014.

[23] A. Sardini, J. S. Amey, K. H. Weylandt, M. Nobles, M. A. Valverde, and C. F. Higgins. Cell volume regulation and swelling-activated chloride channels. Biochim. Biophys. Acta., 1618:153–162, 2003.

[24] J. V. Halliwell, T. D. Plant, J. Robbins, and Nick. B. Standen. Voltage clamp techniques. In N. B. Standen, P. T. A. Gray, and M. J. Whitaker, editors, Microelectrode Technique. The Plymouth Workshop Handbook, pages 13–28. The Company of Biologists Limited, Cambridge, 1987.

[25] G. B. Ermentrout and D. H. Terman. Mathematical Foundations of Neuroscience. Springer, New York, NY, 2010.

[26] W. W. Douglas, T. Kanno, and S. R. Sampson. Influence of the ionic environment on the membrane potential of adrenal chromaffin cells and on the depolarizing effect of acetylcholine. J. Physiol., 191:107–121, 1967.

[27] F. Si, B. Li, W. Margolin, and S. X. Sun. Bacterial growth and form under mechanical compression. Sci. Rep., 5:11367, 2015.

[28] Y. Li, Y. Mori, and S. X. Sun. Flow-driven cell migration under external electric fields. Phys. Rev. Lett., 115:268101, 2015.

[29] J. M. Russell. Sodium-Potassium-Chloride cotransport. Physiol. Rev., 80(1):211–276, 2000.

[30] M. R. Bennett, L Farnell, and W. G. Gibson. A quantitative model of cortical spreading depression due to purinergic and gap-junction transmission in astrocyte networks. Biophys. J., 95(12):5648–5660, 2008.

[31] D. C. Gadsby, J. Kimura, and A. Noma. Voltage dependence of Na/K pump current in isolated heart cells. Nature, 315:63–65, 1985.

[32] J. Gao, R. T. Mathias, I. S. Cohen, and G. J. Baldo. Two functionally different Na/K pumps in cardiac ventricular myocytes. J. Gen. Physiol., 106(5):995–1030, 1995.

[33] A. Bueno-Orovio, C. Sánchez, E. Pueyo, and B. Rodriguez. Na/K pump regulation of cardiac repolarization: insights from a systems biology approach. Pflugers Arch., 466(2):183–193, 2014.

[34] C. M. Armstrong. The Na/K pump, Cl ion, and osmotic stabilization of cells. Proc. Natl. Acad. Sci. U.S.A, 100(10):6257–6262, 2003.

[35] H. Lodish, A. Berk, P. Matsudaira, C. A. Kaiser, M. Krieger, M. P. Scott, L. Zipursky, and J. Darnell. Molecular Cell Biology.

[36] K. Zierler, E. M. Rogus, R. W. Scherer, and F.-S. Wu. Insulin action on membrane potential and glucose uptake: effects of high potassium. Am. J. Physiol., 249(1):E17–E25, 1985.

[37] D. M. Berman, C. Pena-Rasgado, and H. Rasgado-Flores. Changes in membrane potential associated with cell swelling and regulatory volume decrease in barnacle muscle cells. J. Exp. Zool., 268(2):97–103, 1994.

[38] A. Y. Mulkidjanian, A. Y. Bychkov, D. V. Dibrova, M. Y. Galperin, and E. V. Koonin. Origin of first cells at terrestrial, anoxic geothermal fields. Proc. Natl. Acad. Sci. U.S.A., 109(14):E821–E830, 2012.

[39] S. F. Pedersen, E. K. Hoffmann, and J. W. Mills. The cytoskeleton and cell volume regulation. Comp. Biochem. Physiol. A Mol. Integr. Physiol., 130:385–399, 2001.

[40] L. He, J. Tao, F. Si, Y. Wu, T. Wu, V. Prasath, D. Wirtz, and S. X. Sun. Role of membrane-tension gated ca flux in cell mechanosensation. J. Cell Sci., page 134395, 2017.

